# Engineering *E. coli* for magnetic control and the spatial localization of functions

**DOI:** 10.1101/2020.01.06.895623

**Authors:** Mary Aubry, Wei-An Wang, Yohan Guyodo, Eugénia Delacou, Jean Michel Guignier, Olivier Espeli, Alice Lebreton, François Guyot, Zoher Gueroui

## Abstract

The fast-developing field of synthetic biology enables broad applications of programmed microorganisms including the development of whole-cell biosensors, delivery vehicles for therapeutics, or diagnostic agents. However, the lack of spatial control required for localizing microbial functions could limit their use and induce their dilution leading to ineffective action or dissemination. To overcome this limitation, the integration of magnetic properties into living systems enables a contact-less and orthogonal method for spatiotemporal control. Here, we generated a magnetic-sensing *Escherichia coli* by driving the formation of iron-rich bodies into bacteria. We found that these bacteria could be spatially controlled by magnetic forces and sustained cell growth and division, by transmitting asymmetrically their magnetic properties to one daughter cell. We combined the spatial control of bacteria with genetically encoded-adhesion properties to achieve the magnetic capture of specific target bacteria as well as the spatial modulation of human cell invasions.

## Introduction

By programming and harnessing the cellular behavior of living organisms, synthetic biology tools enable broad applications ranging from basic biology to health and environment issues. Synthetic circuits have been developed for *in vitro* and *in vivo* diagnostics^1^, to produce novel material^2,3^, or to direct the assembly of synthetic multicellular systems^4,5^. For instance, programmed as whole-cell biosensors bacteria can report on environmental changes, detect specific molecules^6,7^, or monitor and diagnostic diseases^8–13^. Bacteria can be further modified to act on their environment as illustrated by their use to target pathogenic bacteria^14,15^ or cancer cells^16–19^.

Programming cells to be sensitive to non-biochemical stimuli, such as acoustic or magnetic waves, could expand their capacity to probe or act on their environment^20^. For instance, the integration of magnetic properties into living organisms could enable their spatial manipulation by magnetic forces, and their use as contrast agents for magnetic resonance imaging or as heat generator^21–27^. As future perspectives, the magnetic localization of programmed bacteria may overcome their spatial dissemination driving to ineffective action or to biosafety issues. In this context, several strategies have been established to produce and use magnetic-sensing bacteria. First, magnetotactic bacteria are among the few living systems known to exploit magnetism by using their unique intracellular organelles, the magnetosomes^28^, to swim along the Earth’s magnetic field. Despite several attempts to use magnetotactic bacteria^29^, they remained difficult to harness^30^ and to manipulate genetically. One second strategy consisted in building bacterial biohybrid systems either using magnetotactic bacteria carrying cargo-particles^31,32^, or reciprocally, using a magnetic field to control the orientation of motile bacteria linked to magnetic beads^33,34^. A third approach aimed to magnetize naturally diamagnetic microorganisms or eukaryotic cells by over-expressing iron-storage ferritins or iron-binding proteins inside their cytoplasm. These bacteria could serve as containers favoring the formation of iron-oxide deposits when cells were fed with iron^35–40^. These studies showed that mineralized cells contained iron oxide deposits, can be detected using NMR, and can be magnetically sorted. However, to envision biotechnological applications using mineralized cells several important challenges still need to be achieved. Among primary questions, knowing how magnetic properties are transmitted during cell division, or whether magnetized cells are amenable for achieving defined biochemical functions while being magnetically manipulated, are essential elements that have not yet been solved.

To address such questions, we engineered and characterized *MagEcoli*, that are iron-mineralized *Escherichia coli* bacteria expressing the iron-storage ferritin. We used *MagEcoli* to demonstrate that mineralized bacteria can be programmed to perform specific biochemical functions with spatiotemporal control using magnetic forces. First, we performed structural and magnetic characterization of *MagEcoli* and found that they contained iron oxide-enriched bodies conferring magnetic properties. Next, we showed that *MagEcoli* could be spatially manipulated when exposed to magnetic forces, with an efficiency that increased with iron loading. Moreover, *MagEcoli* divided and transmitted asymmetrically iron oxide ferritin-enriched bodies during division, thus avoiding the dilution of the magnetic properties during population growth. Finally, we combined the spatial control of *MagEcoli*, modified with genetically encoded-adhesion properties displayed on their outer membrane, to achieve the magnetic capture of specific target cells as well as the spatial modulation of human cell invasions.

## Results

### Genetic and chemical modifications to obtain a magnetic *Escherichia Coli*

We aimed to direct the formation of iron-bearing within *Escherichia Coli* cytoplasm to provide magnetic properties to the bacteria. Our strategy to produce iron-rich inclusions in bacteria relied on a two-step process consisting first in overexpressing fluorescently-labelled ferritins and then supplying Fe(II) to the growth medium to biomineralize the bacteria. We chose the heterologous production of the iron-storage ferritins derived from *Pyroccocus Furiosus*^41^. After 16 hours of iron biomineralization, bacteria were washed and then characterized at the nanometer scale using transmission electron microscopy (TEM) images of cross sectioned mineralized *E. coli*. TEM images showed accumulation of a large electron-dense-deposit often localized at the extremity of the bacteria (**Fig. 1A**). The intracellular clusters were formed by the aggregation of small nanoparticles (~ 3-5 nm, **Fig. 1B**), which was consistent with the cavity size of ferritin nanocages (8 nm inner diameter). The iron-rich clusters, quasi-spherical in shape and 100-300 nm in diameter, were localized in the cytosol of the bacteria. The electron diffraction pattern of the iron clusters showed that the nanoparticles were either amorphous or poorly crystallized (**Fig. 1D**). Their analysis by energy dispersive X-ray spectroscopy in scanning transmission electron microscopy (STEM) on 60 nm thick cross-sections of *Escherichia Coli* overexpressing ferritin proteins revealed iron, phosphorous and oxygen (**Fig. 1C, E-H**). This was confirmed by analyzing the minerals inside entire bacteria using cryo-TEM (**Fig. S1A**). As control, we imaged mineralized GFP-expressing *E. coli* which did not overexpressed ferritins, and no-electron dense deposits were observed in those bacteria (**Fig. S1B**). In those condition, only extra cellular precipitates were observed suggesting that the overexpression of ferritins was necessary to induce the formation of intracellular iron oxide nanoparticles (**Fig. S1C**).

**Figure 1:**
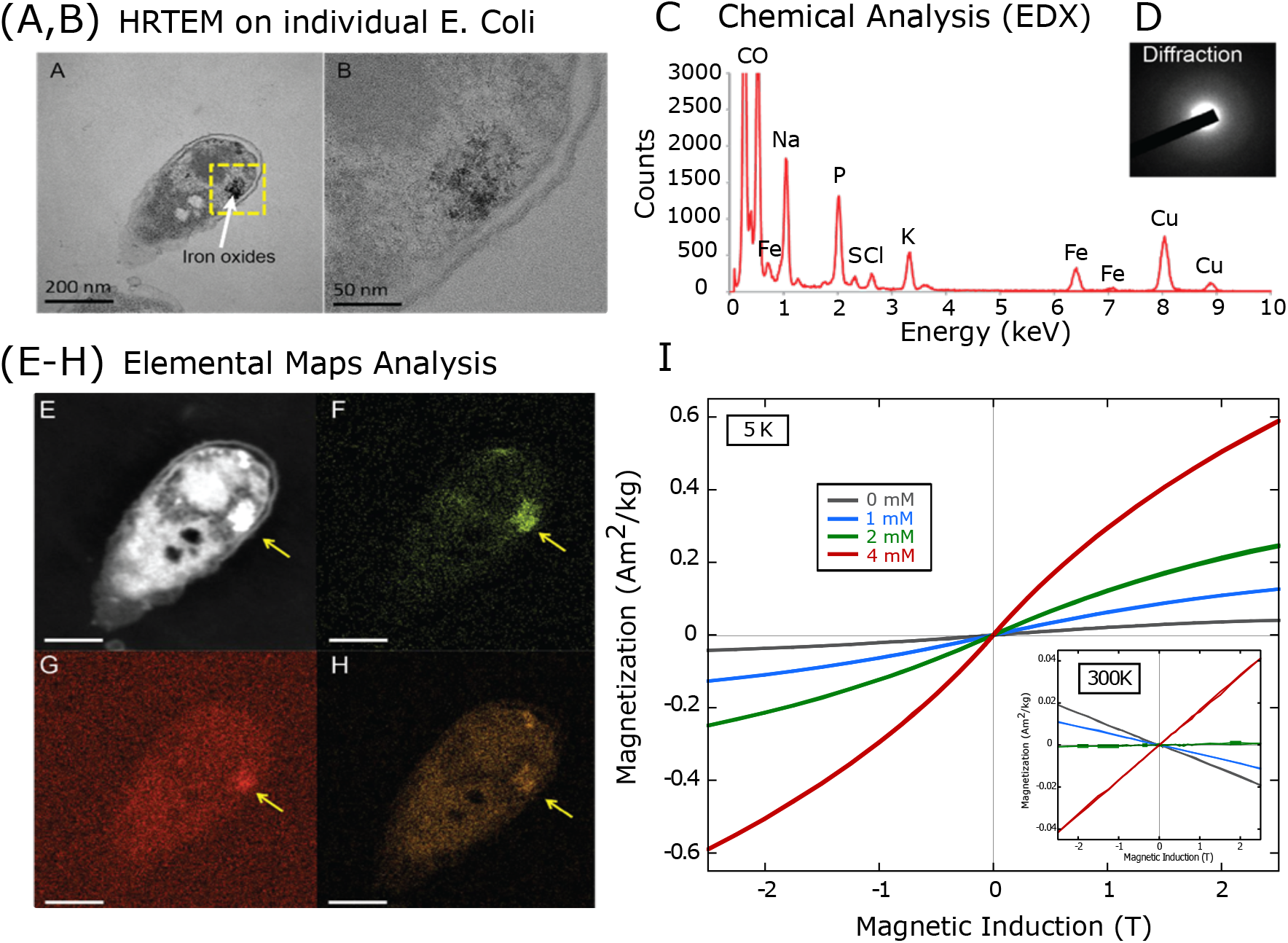
Structural and chemical characterization of *MagEcoli* bacteria. **(A,B)** TEM images of a cross sectioned mineralized mCherry-ferritin expressing *E. coli* strain. **(C)** Energy dispersive X-ray spectroscopy spectra of the electron - dense deposit (yellow area). **(D)** Electron diffraction pattern of nanoparticles. **(E - H)** Elemental mapping of a mineralized cross-section of mCherry-ferritin expressing *E. coli* strain. **(E)** STEM image. **(F-H)** Each panel represents the detection of a different element: Iron (F), Oxygen (G), Phosphorus (H). Scale bar, 400 nm. **(I)** Magnetization curves of mineralized *E. coli* for different concentration of iron supplementation: 0 mM (black), 1 mM (blue), 2 mM (green), 4 mM (red). Measurements were performed at 5 K and 300K (insert) on a MPMS, with magnetic inductions cycling between +2.5T, −2.5T, and +2.5T.

Further quantitative characterization of the bacteria magnetic properties was achieved through the use of a magnetic properties measurement system (MPMS). The cells were subjected to a measurement of their mass-normalized magnetization at 5 and 300 Kelvin in magnetic inductions ranging between −2.5 and 2.5T (**Fig. 1I**). At room temperature (*i.e*., 300K), all samples exhibit a linear magnetization-versus-field behavior. The slope of the magnetization curve is negative for the sample without iron supplementation, which is inherent to the diamagnetic nature of most biological materials. Iron supplementation resulted in the addition of another linear component of positive slope, likely of paramagnetic nature. The maximum magnetization (in 2.5T) was increased by 0.01, 0.02 and 0.06 Am^2^/kg for the 1, 2, and 4 mM Fe supplementations, respectively. The gain of magnetic susceptibility due to iron biomineralization is more evident when measuring at low temperature, as paramagnetism and other magnetic properties (ferro-, ferri-, antiferro-magnetism) increase in magnitude as temperature decreases, while diamagnetism remains constant. Measurements preformed at 5K clearly display this magnetic enhancement. Compared to the zero-supplementation sample, maximum values of the magnetization increased by 0.09, 0.21, and 0.55 Am^2^/kg for the 1, 2, and 4 mM Fe supplementations, respectively. Altogether, those data highlight that magnetic *E. coli* contain iron minerals ferritin-enriched bodies conferring magnetic properties (referred as *MagEcoli* here after).

### Spatial manipulation and localization of bacteria upon magnetic forces

To assess the possibility to spatially manipulate *MagEcoli*, we performed magnetophoretic experiments, which consist in observing the motion of non-motile bacteria submitted to magnetic forces. A mixture of biomineralized bacteria expressing mCherry-ferritin (*MagEcoli^mCherry^*) and non-mineralized ones, expressing emGFP-ferritin (*E. coli^GFP^*) were diluted in a minimal medium with a density adjusted to prevent bacterial sedimentation. The mixture was then confined into water-in-oil droplets to minimize hydrodynamic flow perturbations and facilitate observation. Once formed, the bacteria droplets were injected into a capillary next to a permanent magnet generating a gradient of about 10 T.m^−1^. Time-lapse observations showed that within few minutes the *MagEcoli^mCherry^* began to move in a direction oriented towards the magnet, whereas non-mineralized ones displayed no net motion (**Fig. 2A**). Moving bacteria eventually accumulated on the edge of the droplet as illustrated by the strong enhancement of mCherry signal intensity (**Fig. 2A**). During this process, *E. coli^GFP^* remained uniformly distributed within the droplet. After 180 minutes almost all magnetic bacteria were attracted (**Fig. 2A**). In order to quantify the mobility of the *MagEcoli*, we tracked single bacterial trajectories within the droplet and computed their speed (**Fig. 2B**). This procedure was performed for respectively 1, 2, 3, and 4 mM Fe(II) added during the biomineralization step. For instance, single bacteria that were mineralized with 4mM iron displayed a directed motion towards the magnet position with a mean speed of about 5.3+/-1.6 μm/min (mean +/- standard deviation) (**Fig. 2C**). These mean speed values were also strongly correlated to the magnetic enhancement values deduced from MPMS measurements. This asymmetrical magnetic concentration procedure can be applied to force the co-localization of two bacterial populations, as exemplified in **Fig. S2** where *MagEcoli^GFP^* and *MagEcoli^mCherry^* were strongly concentrated within the same area at the vicinity of the magnet.

**Figure 2:**
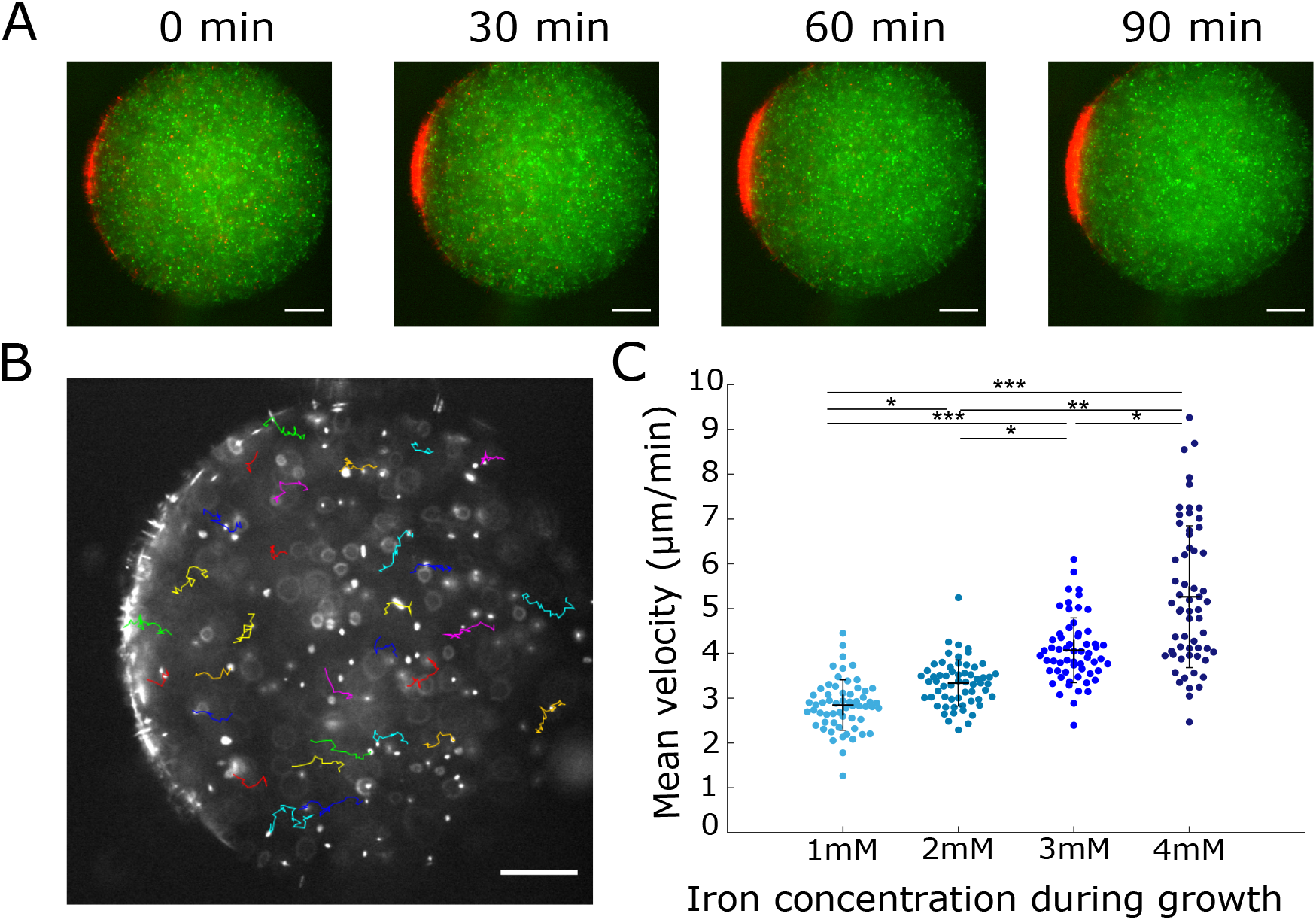
Biomineralized bacteria can be micromanipulated through space with a magnetic field. **(A)**Representative time lapse epifluorescence acquisition of the magnetic localization of *MagEcoli^mCherry^* in a confined environment upon magnetic force application. *MagEcoli^mCherry^* were homogenously mixed with nonmagnetic *E.coli^GFP^* at early time point. The magnet was positioned on the left. Time points at 0, 30, 60, 90 min after starting acquisition. Scale bar, 60 μm, color merged. **(B)** Representation of trajectories as a function of time of *MagEcoli^mCherry^* mineralized with 2 mM of iron II. Magnet on the left. Scale bar, 60μm. **(C)** Histogram of bacterial speed during magnetophoresis experiments as a function of iron concentration during biomineralization. For each condition, the mean +/- standard deviation are displayed, for two independent experiments. NS means there is no significant difference between the two distributions, one star means p-value < 10^−5^, two stars mean p-value < 10^−10^, three stars mean p-value < 10^−15^.

Altogether, these data showed that *MagEcoli* can be spatially manipulated upon magnetic forces, with an efficiency that increases with the concentration of iron added during the biomineralization step. The magnetic concentration process is very specific of the state of biomineralization of the bacteria and did not affect non-magnetized bacteria diffusing in the mixture, allowing to perform basic operations such as magnetic separation and magnetic mixing (**Fig. 2A and Fig. S2**).

### Becoming of *MagEcoli* over time: How magnetic properties propagate through cell division?

Obtaining magnetized bacteria that can sustain cell division is of primary importance for basic understanding, and also to envision applications requiring magnetic manipulations of metabolically active bacteria as well as long term operations. We examined the transmission of magnetic properties after cell division by combining microscopy observations and magnetophoresis.

First, after over-night mineralization, *MagEcoli^mCherry^* were diluted and let grow into fresh LB medium lacking iron supply. We observed ferritin-enriched bodies within bacteria at various growth stages: before new growth, and after 1, 2 and 3 divisions respectively. As the bacteria grew in absence of IPTG, mCherry fluorescence was directly correlated with the presence of ferritins expressed by the mother bacteria, which allowed us to monitor the becoming of iron oxide enriched bodies (**Fig. 3A**). Before division, almost 100% of bacteria displayed heterogeneous mCherry fluorescence accumulation (**Fig. 3A**). At this stage, we found bright fluorescent bodies localized at one or at both bacterial poles and coexisting with a diffuse fluorescence distributed within the cell body. These observations were in agreement with TEM acquisitions (**Fig. 1A**). When observing bacteria at a later stage of growth, the number of fluorescent bacteria decreased compared to non-fluorescent ones. After the 1^st^ division, 30% of the observed bacteria continued to display a fluorescent accumulation, whereas in contrast, the remaining bacteria showed a weak or no fluorescence signal (**Fig. 3A**). The fluorescent bacteria represent about 10% of the total bacteria after the third division, suggesting that bacteria asymmetrically transmitted their ferritin-enriched bodies to daughter cells (**Fig. 3A**).

**Figure 3:**
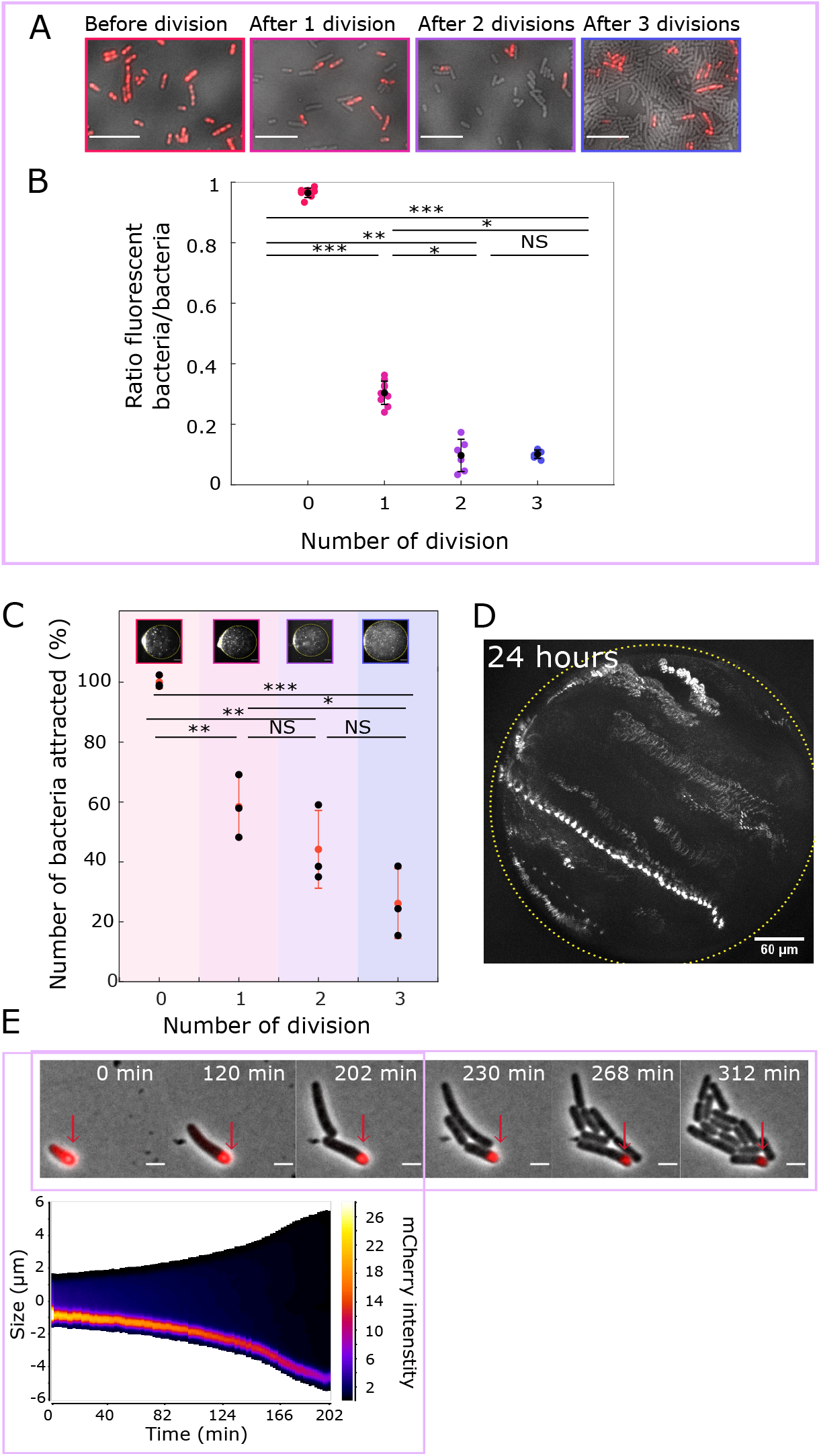
Evolution of the magnetic properties of *MagEcoli* as a function of cell division. **(A)** Superimposition of mCherry fluorescence and bright field images of mineralized bacteria after 0, 1, 2, and 3 divisions. Scale bar, 10 μm. **(B)** Quantification of the evolution as a function of cell division of the ratio of bacteria remaining magnetic compared to the total number of growing bacteria (1000 bacteria, two different experiments performed at different days). Each point represents a ratio computed from a microscopic observation. The mean +/- standard deviation are displayed. NS means there is no significant difference between the two distributions, one star means p-value < 10^−5^, two stars mean p-value < 10^−10^, three stars mean p-value < 10^−15^. **(C) Upper panel:** Representative magnetophoresis images of the accumulation of *MagEcoli* after 0, 1, 2, and 3 divisions. Images were taken 90 min after starting the accumulation; magnet is on the left, scale bar, 60 μm. **Graph:** Quantification of the number of bacteria attracted towards the magnet for the corresponding experiment. For each condition, the mean +/- standard deviation is displayed for three independent experiments. NS means there is no significant difference between the two distributions, one star means p-value < 0.1, two stars mean p-value < 0.01, three stars mean p-value < 0.001. **(D)** Projection of bacterial trajectories integrated on 50 minutes of bacteria still containing mineralized ferritins after 24 hours of new growth. Magnet on the bottom left. Scale bar, 60 μm. **(E)** Time-lapse images of live fluorescence microscopy images (Merged images of phase contrast and mCherry channels, level of mCherry adjusted for each image). Time points at 0, 120, 202, 230, 268 and 312 min. Scale bar, 2 μm. **Lower panel:** Kymograph of the dividing bacteria displayed on top.

This asymmetric cell division model was also confirmed by monitoring the process of division using live microscopy (**Fig. 3E**). *MagEcoli^mCherry^* that divided after biomineralization conserved their large bright inclusion bodies at the pole, leading to the transmission of the major part of mCherry-ferritin to one daughter cell only.

Furthermore, after 24 hours of growth following the end of biomineralization, the remaining fluorescent bacteria moved as fast as the mineralized bacteria that had not undergone cell division (4.9 μm.min^−1^ +/- 3.1 μm.min^−1^, 15 tracked trajectories, 3 different assays, (**Fig. 3D**)). We next quantified the evolution as a function of cell division of the ratio of bacteria remaining magnetic compared to the total number of growing bacteria. Mineralized mother bacteria were grown in a medium supplemented with IPTG to allow all newborn bacteria (magnetic and non-magnetic) to be monitored by fluorescence. The ratio of attraction towards the magnet of magnetic versus nonmagnetic bacteria drop from 100 to 58, 44, and, 26% after the first three division steps (**Fig. 3C**). Altogether, these data showed that *MagEcoli* were still able to grow and divide. Only newborn bacteria maintaining iron oxide ferritin-enriched bodies inherited magnetic properties. This asymmetric division process avoids the dilution of the magnetic properties during population growth.

### *MagEcoli* with genetically encoded-adhesion properties for spatial control of cell capture and cell invasion

Engineering the adhesion properties of cells offers multiple applications ranging from programming tissues, living materials, cell signaling, as well as designing whole-cell biosensors to detect specific analytes^2–6,16,18^. To envision applications combining the spatial control of bacteria and adhesion, we extended the capacity of *MagEcoli* to perform two distinct specific functions: the capture of specific bacteria and the invasion of human cells.

In order to capture, manipulate, or sort in space specific target bacteria, we have implemented in *MagEcoli* a genetically encoded surface-displaying adhesin system developed for controlling cell-cell adhesion^5^. This modular system displays on bacteria outer membrane nanobodies or antigens^5^. *MagEcoli* were transformed to produce on their outer membrane Ag2, an antigen based on a cell surface-bound adhesin and encoded as a single fusion protein designed to specifically recognized Nb2 nanobody-presenting bacteria^5^ (**Fig. 4A**). Expression of Nb2 and Ag2 was under control of anhydrotetracycline addition. When Ag2-producing *MagEcoli MagEcoli^Ag2/mCherry^*) were mixed with Nb2-producing bacteria (*E.coli^Nb2/GFP^*), we could observe multicellular aggregates formed by few tens of cells with the same morphological patterns as previously demonstrated (**Fig. 4B, left panel and S3**). This indicated that the mineralization of bacteria did not preclude their adhesive properties. In absence of anhydrotetracycline, no aggregation was observed confirming the specificity of the adhesion system (**Fig. 4B, right panel**). To assess the capacity to capture and spatially manipulate target bacteria using *MagEcoli*, we mixed *MagEcoli^Ag2/mCherry^* with *E. coli^Nb2/GFP^* in droplets and applied a permanent magnetic field as explained above (**Fig. 4A**). Remarkably, at the vicinity of the magnet, the concentration of *E.coli^Nb2/GFP^* was observed concomitantly with the one of *MagEcoli^Ag2/mCherry^* (**Fig. 4C and 4D**). After about 30 min of attraction, we observed an increase in GFP as well as mCherry intensity at the vicinity of the magnet, indicating that *E. coli^Nb2/GFP^* were dragged along the magnetic gradient by the *MagEcoli^Ag2/mCherry^*. Moreover, no attraction was observed in the absence of anhydrotetracycline and the attraction of *MagEcoli^Ag2/mCherry^* left the position of *E.coli^Nb2/GFP^* unaltered (**Fig. 4D**). With a closer look at the magnetophoresis experiments, we observed long-range transport of *E. coli^Nb2/GFP^* by *MagEcoli^Ag2/mCherry^* along magnetic force axis, indicating that enrichment by target cells was powered by *MagEcoli* transportation (**Fig. 4E**). Altogether these data showed that *MagEcoli* can be programmed to capture and transport specific bacteria upon magnetic field application.

**Figure 4:**
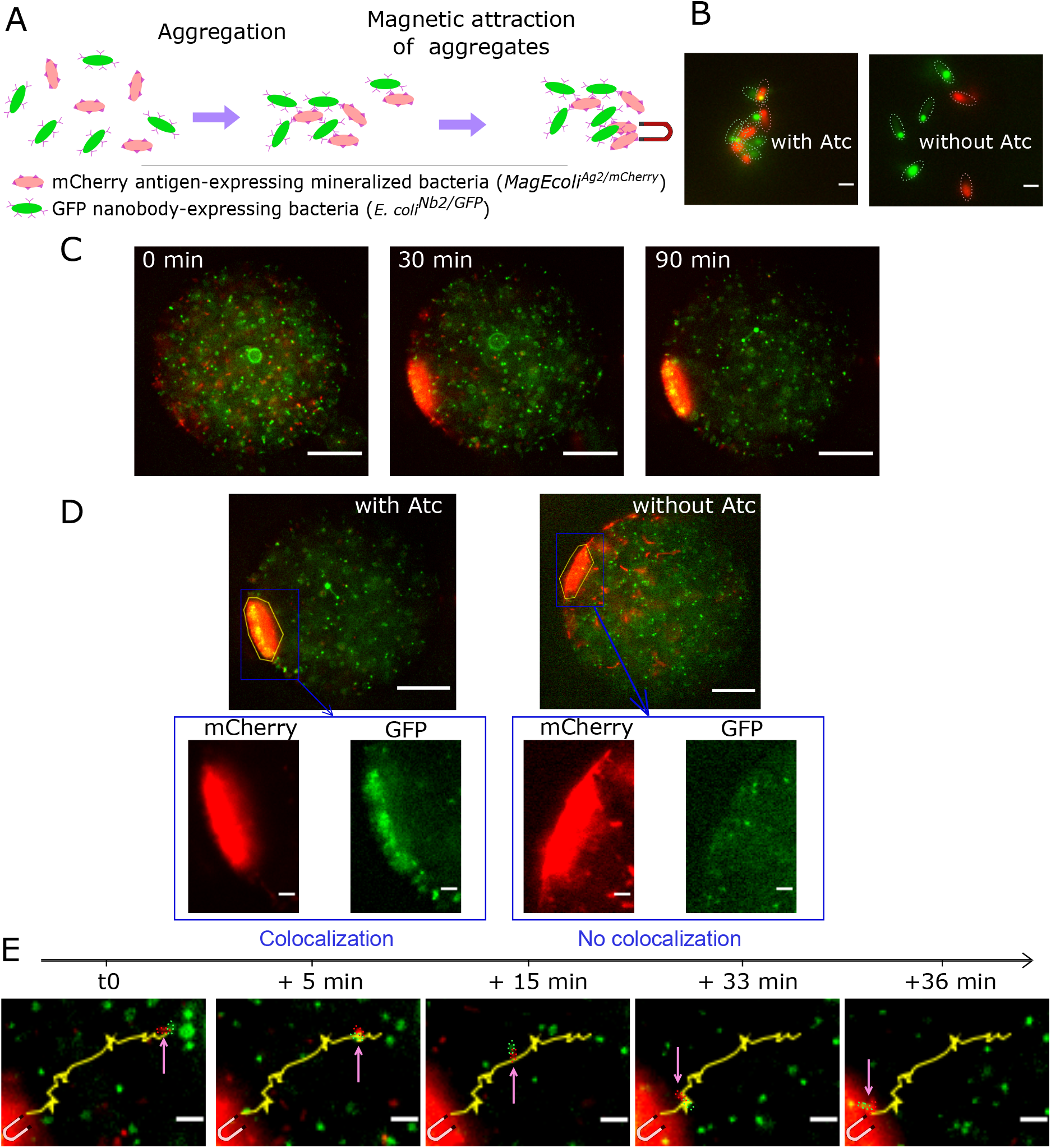
Capture and spatial attraction of targeted bacteria by antigen/antibody recognition. **(A)** Scheme of the assay of the capture and spatial attraction of targeted bacteria by antigen/antibody recognition. GFP nanobody-expressing bacteria (*E. coli^Nb2/GFP^*) can adhere to mCherry antigen-expressing mineralized *E. coli*(*MagEcoli^Ag2/mCherry^*). **(B)** On the **left panel**: aggregation of antigen-presenting *MagEcoli* (mCherry) with nanobodypresenting *E. coli* (GFP), in presence of anhydrotetracycline (Atc). On the **right panel**: control performed without anhydrotetracycline. Epifluorescence observations. Merged images. Scale bar, 2 μm. **(C)** Time lapse images of magnetic accumulation of antigen-producing *MagEcoli* (mCherry) adhering to nanobody-producing *E. coli* (GFP). Images at 0 min, 30 min and 90 min upon magnetic field application. Merged images. Scale bar, 60 μm. **(D)** Images of magnetic accumulation of aggregates in the presence (left) or in the absence (right) of anhydrotetracycline. Merged images. Scale bar 60μm. **Below**: zoom of the accumulation of magnetic bacteria. Colorized images. Extracted from the movie in panel C. Scale bar, 10 μm. **(E)** Time-lapse images showing the trajectory two adhering bacteria (*MagEcoli^Ag2/mCherry^* and *E. coli^Nb2/GFP^*) over time. Extracted from the movie in panels C and D. Merged images. Scale bar, 10 μm.

We next devised an assay to monitor the spatial localization of *MagEcoli* programmed to invade human cells. First, we expressed in *MagEcoli* the gene encoding *Yersinia pseudotuberculosis* invasin, an adhesive protein that is known to allow the invasion of cultured animal cells by otherwise non-invasive enterobacteria^42^. We verified that *MagEcoli^inv/GFP^*(*MagEcoli* expressing invasin) were able to specifically recognize and invade human cells (Fig. 5A and S4). The invasiveness of *MagEcoli^inv/GFP^* towards HeLa cells was quantified by gentamicin protection assay (Fig. S4): extracellular bacteria are killed by the antibiotic, while intracellular bacteria are protected due to the impermeability of host cells plasma membranes. Plating serial dilutions of cell lysates on LB-agar plates following 1 hour of gentamicin treatment thus enables an estimation of the number of internalized bacteria. We found that the internalization of *MagEcoli^inv/GFP^* into HeLa cells did not differ significantly from that of the same bacteria that had been grown in absence of iron (Fig. S4), arguing that mineralization of invasive *E. coli* did not impair their ability to invade human cells. To demonstrate the magnetic localization of cell invasion, we placed a magnet under a dish containing HeLa cells in contact with *MagEcoli^inv/GFP^* for 4 hours (Fig. 5B). After gentamicin treatment, cells were fixed and stained to image *MagEcoli*, actin filaments, as well as the cell nucleus. Strikingly, cells adhering in the vicinity of the magnetic field contained a larger number of *MagEcoli* in their cytoplasm than cells adhering far from the magnet. Moreover, the density of bacteria per cells was correlated with the proximity with the magnetic field and reach about 8-fold (Fig. 5C and 5D). Altogether, these data demonstrate our ability to target *MagEcoli* invasion to a specific zone with a magnetic field.

**Figure 5:**
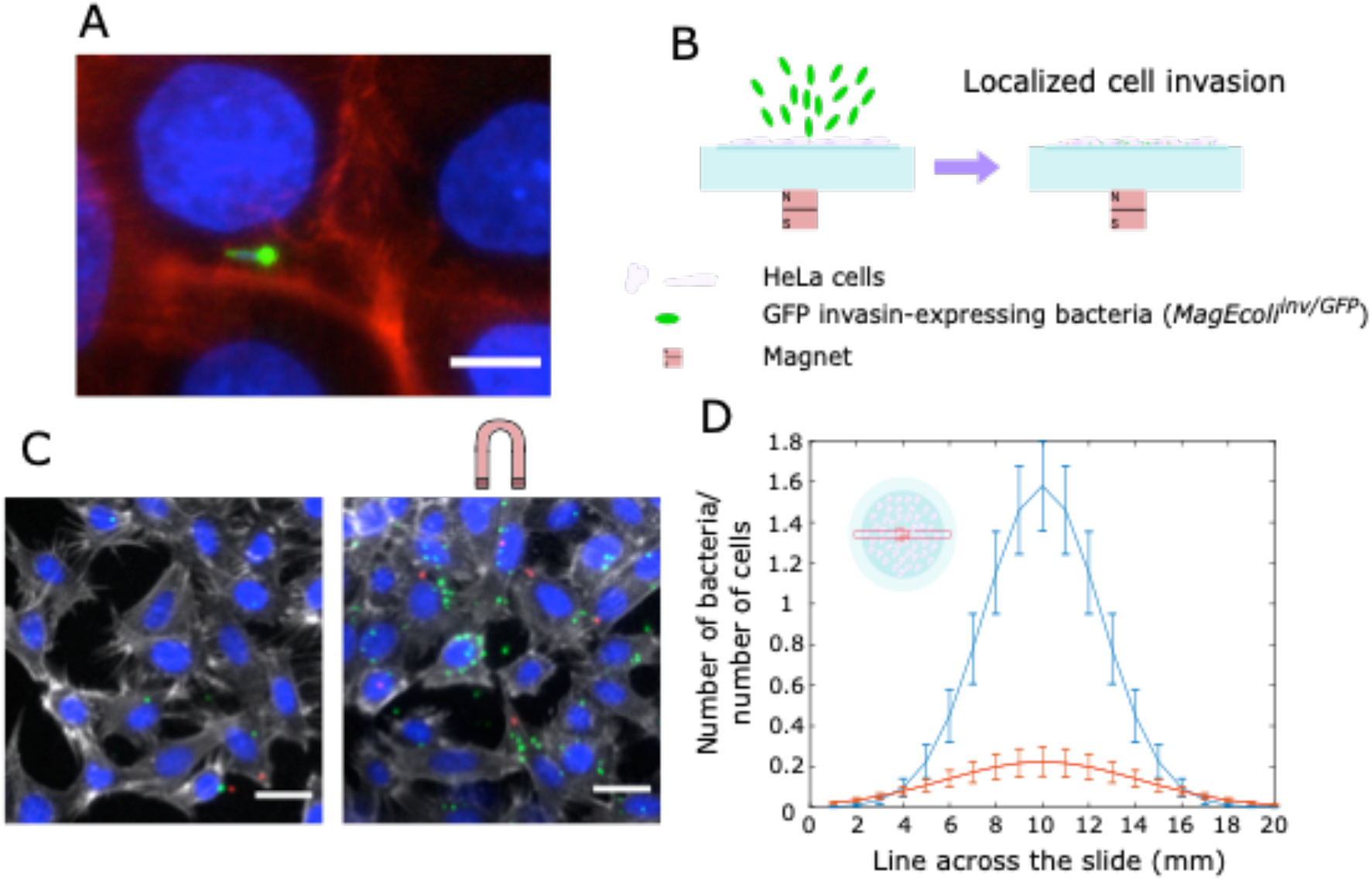
Magnetic localization of bacterial infection of HeLa cells. **(A)** Epifluorescence images of *MagEcoli^inv/GFP^* inside a Lovo cell. In blue, nucleus of human cells (LoVo) and bacterial DNA; in red, actin; in green, ferritin of *MagEcoli^inv/GFP^*. Scale bar, 10 μm. **(B)** Scheme of the set-up used to localize invasion of HeLa Cells by magnetic invasive bacteria. *MagEcoli^inv/GFP^* are placed on a culture dish covered with Hela Cells. **(C)** Epifluorescence images of invasion of Hela cells by *MagEcoli^inv/GFP^*. In blue, nucleus of HeLa cells; in gray, actin; in green, *MagEcoli^inv/GFP^*; in red, *E. coli^inv/mCherry^*. Scale bar, 30 μm. **(D)** Number of bacteria that invaded one HeLa Cell. In blue, the mean of the number of *MagEcoli^inv/GFP^* for 4 different zones of observation on the same sample; in red, the respective number of *E. coli^inv/mCherry^*. The x-axis represents the zone of observation, in mm. Data are normalized by the number of nucleus of cells counted on each field of observation.

## DISCUSSION

We demonstrate the spatial control of engineered bacteria mediated by magnetic forces and programmed to achieve specific tasks using modified surface-adhesion properties. Magnetic bacteria were engineered using a two-step processes consisting in expressing the iron storage ferritin in *E. coli* and growing these bacteria in an iron-riche medium. Iron mineralization of ferritin-expressing bacteria resulted in the formation of amorphous iron oxide minerals enriched in iron, oxygen and phosphorus. These *MagEcoli* bacteria display paramagnetic properties that increase with the amount of iron supplemented during bacteria growth. In contrast, biomineralized bacteria that did not over-express ferritins were not exhibiting any detectable intracellular iron oxide particles, but showed extracellular iron deposits. MPMS measurements showed that biomineralized control *E. coli* displayed a diamagnetic signal, suggesting that the paramagnetic contribution of *MagEcoli* is mainly due to intracellular iron oxide minerals. These data suggest a model in which ferritin-expressing *E. coli* hyperaccumulate iron metals that form iron-oxide minerals stored into ferritin-enriched bodies, with a chemical and crystal-structure that are constrained by the physical-chemistry state of the *E. coli* cytoplasmic in terms of pH/redox conditions.

We next demonstrate that *MagEcoli* can be concentrated using magnetic forces to generate spatial heterogeneity in bacterial concentration. For instance, when confined in a millimeter-size confined environment, *MagEcoli* could be physically separated from a nonmagnetic bacteria population or, in contrast, forced to mix together with a second magnetic bacterial strain in a specific area.

To investigate how cell division could impact the magnetic properties of *MagEcoli*, we studied how ferritin-enriched bodies and magnetic properties propagated when bacteria divided. We found that *MagEcoli* transmitted asymmetrically their ferritin-enriched bodies to only one daughter cell. The proportion of attracted magnetic bacteria by the magnet dropped concomitantly with the cell division number to reach about 10% after three divisions. This corroborates a model for which the main magnetic properties were inherited by one daughter cell, consequently resulting in the maintenance of a constant population of *MagEcoli* in regard to a growing non-magnetic population of bacteria. Interestingly, this mechanism limits the dilution of the magnetic properties, which would in contrast be expected if mineralized ferritin bodies would equally be distributed between the two daughter cells.

One essential aspect when envisioning applications of *MagEcoli* was to assess how these bacteria could be further programmed to perform specific biochemical functions. By expressing genetically encoded surface adhesion proteins, we studied the spatial localization of *MagEcoli* programmed to recognize and adhere to specific bacteria or to invade a mammalian host. We first demonstrated the spatial manipulation of *MagEcoli* acting as a surface-displaying antigen to adhere specifically to non-magnetic nanobody-displaying bacteria. We found that *MagEcoli* can capture and transport target bacteria along a magnetic force axis to eventually drive their accumulation. Such properties illustrate potential perspectives for biotechnological purposes as *MagEcoli* could be programmed and magnetically manipulated to transport cargos, for patterning living materials, or as whole-cell biosensors for *in vitro* or *in vivo* diagnostic. In a second application, we demonstrated the spatial modulation of cell invasion in a magnetic field by programming *MagEcoli* to invade human cells. The magneto-localization of infection could provide a novel tool for basic studies of host-pathogen interactions. One horizon consists in the spatiotemporal control of bacteria programmed as delivery vehicles to release cytotoxic molecules into cancer cells.

## Author’s contributions

M.A, F.G, and Z.G conceived the project and analysed results of the study. M.A carried out and analysed most of the experiments. WA.W contributed to the initial design of the magnetophoresis experiments. WA.W, J.M Guignier and F.G carried out and analysed the TEM and energy dispersive X-ray spectroscopy experiments. Y.G carried out and analysed the MPMS data. M.A and O.E carried out live microscopy of bacteria division. A.L carried out the assessment of cell invasion by gentamicin protection assay. M.A and A.L performed the spatial modulation of cell invasion. M.A and E.D performed the magnetic capture assay. M.A and Z.G wrote the manuscript and all authors were involved in revising it critically for important intellectual content.

## Acknowledgements

The authors acknowledge the members of the Biophysical Chemistry group of the École normale supérieure, Ludovic Jullien, and Guillaume Morin for fruitful discussion. This work was supported by the ANR (ANR-16-CE09-0002-Nanoheaters), the CNRS, and Ecole Normale Supérieure.

