## Supplementary Information for "Engineering *E. coli* for magnetic control and the spatial localization of functions"

5- Institut de biologie de l'ENS (IBENS), Département de biologie, École normale supérieure, CNRS, INSERM, PSL University, 75005 Paris, France.

6- INRAE, IBENS, 75005 Paris, France.

7- Institut Universitaire de France (IUF).

**Figure S1.**

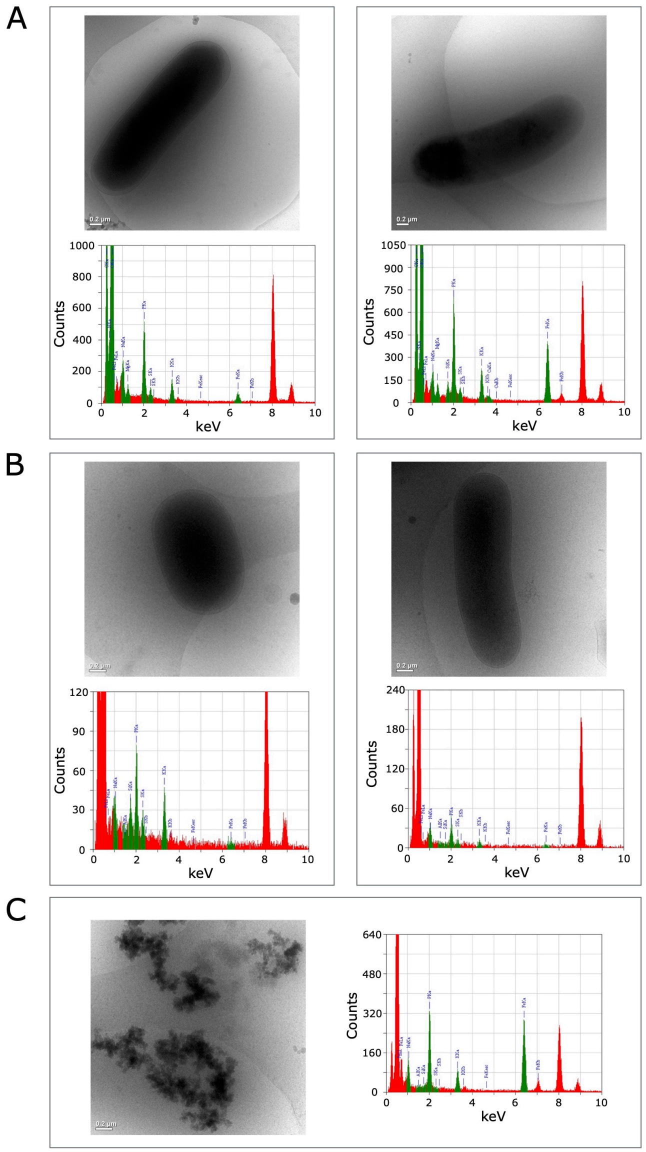


**Figure S1: Cryo-TEM observations of *MagEcoli* and *E. coli* bacteria after biomineralization. (A)** *MagEcoli* mineralized with 2mM Fe(II) (**Right panel**) or 4 mM Fe(II) (**Left panel**). On the **top panel**: high resolution cryo-TEM images. Scale bar, 0.2µm. On the **bottom panel**: energy dispersive X-ray spectroscopy spectra for the bacterial cytoplasm. **(B)** Control *E. coli* that does not overproduce ferritin in presence of 2mM Fe(II) (**Right panel**) or 4 mM Fe(II) (**Left panel**). On the **top panel**: high resolution cryo-TEM images. Scale bar, 0.2µm. On the **bottom panel**: energy dispersive X-ray spectroscopy spectra for the bacterial cytoplasm. **(C)** Extracellular aggregates generated in the medium of control *E. coli* that does not overproduce ferritin in the presence of 4 mM Fe(II). On the **top panel**: high resolution cryo-TEM images. Scale bar, 0.2µm. On the **bottom panel**: energy dispersive X-ray spectroscopy spectra for the aggregate.

**Figure S2.**

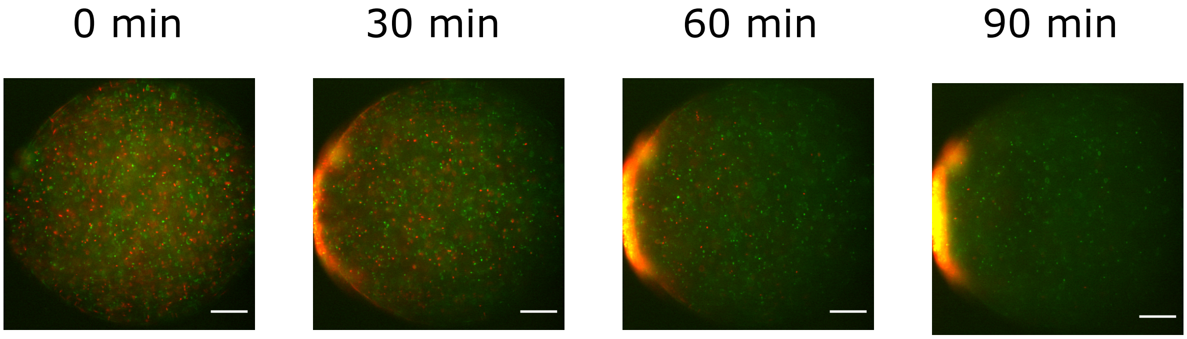


**Figure S2: Representative time lapse fluorescence acquisition of the magnetic localization of *MagEcoli*$^{mCherry}$ and *MagEcoli*$^{GFP}$ in a confined environment upon magnetic force application.** *MagEcoli*$^{mCherry}$ were homogenously mixed with *MagEcoli*$^{GFP}$ at early time point. The magnet was positioned on the left. Time points at 0, 30, 60, 90 min after starting acquisition. Scale bar, 60 µm, color merged.

**Figure S3.**

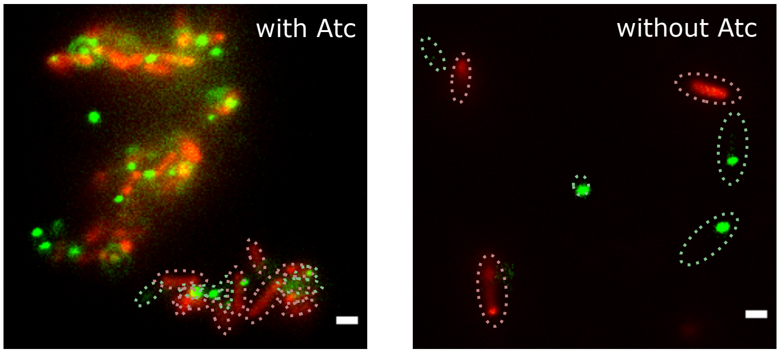


**Figure S3: Fluorescence images of aggregates of *MagEcoli*$^{Ag2/mCherry}$ with *E. coli*$^{Nb2/GFP}$.** On the **left panel**: aggregates of *MagEcoli*$^{Ag2/mCherry}$ and *E. coli*$^{Nb2/GFP}$ in presence of anhydrotetracycline (Atc). On the **right panel**: control performed without anhydrotetracycline. Fluorescence observations. Merged images. Scale bar, 2 µm.

**Figure S4.**

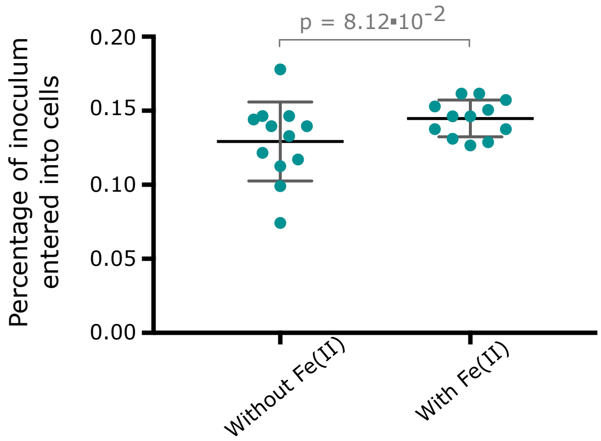


**Figure S4: The biomineralization of invasive *MagEcoli* does not impede their internalization into epithelial cells.** LoVo cells were infected for 4 h with *E. coli*$^{inv/GFP}$ expressing the invasin gene from *Y. pseudotuberculosis* and the ferritin-GFP fusion, that had been grown in medium containing either no Fe(II) (*E. coli*$^{inv/GFP}$*, left*) or 4 mM of Fe(II) (*MagEcoli*$^{inv/GFP}$ , *right*). After gentamicin treatment for 1 h, the percentage of the inoculum having entered was assessed by plating serial dilutions of cell lysates. The values obtained for 12 technical replicates from a representative invasion assays among three independent experiments, their means and standard deviations were plotted. The probability *p* of rejection of the null hypothesis was assessed using a two-tailed Student's *t*-test.
